# DUSP1 regulates apoptosis and cell migration, but not the JIP1-protected cytokine response, during Respiratory Syncytial Virus and Sendai Virus infection

**DOI:** 10.1101/163360

**Authors:** Alexa C. Robitaille, Elise Caron, Nicolas Zucchini, Espérance Mukawera, Damien Adam, Mélissa K. Mariani, Anaïs Gélinas, Audray Fortin, Emmanuelle Brochiero, Nathalie Grandvaux

## Abstract

The host antiviral response involves the induction of interferons and proinflammatory cytokines, but also the activation of cell death pathways, including apoptosis, to limit viral replication and spreading. This host defense is strictly regulated to eliminate the infection while limiting tissue damage that is associated with virus pathogenesis. Post-translational modifications, most notably phosphorylation, are key regulators of the antiviral defense implying an important role of protein phosphatases. Here, we investigated the role of the dual-specificity phosphatase 1 (DUSP1) in the host defense against human respiratory syncytial virus (RSV), a pathogenic virus of the *Pneumoviridae* family, and Sendai virus (SeV), a model virus being developed as a vector for anti-RSV vaccine. We found that DUSP1 is upregulated before being subjected to proteasomal degradation. DUSP1 does not inhibit the antiviral response, but negatively regulates virus-induced JNK/p38 MAPK phosphorylation. Interaction with the JNK-interacting protein 1 scaffold protein prevents dephosphorylation of JNK by DUSP1, likely explaining that AP-1 activation and downstream cytokine production are protected from DUSP1 inhibition. Importantly, DUSP1 promotes SeV-induced apoptosis and suppresses cell migration in RSV-infected cells. Collectively, our data unveil a previously unrecognized selective role of DUSP1 in the regulation of tissue damage and repair during infections by RSV and SeV.

## INTRODUCTION

Respiratory syncytial virus (RSV) belongs to the *Pneumoviridae* family of large enveloped negative-sense ssRNA viruses that includes important human pathogens ^1,2^. RSV is a leading cause of acute lower respiratory tract infections associated with morbidity and mortality in infants, children and elderly, but also adults of all-age with a compromised immune system, cardiopulmonary diseases or following transplantation ^3,4^. The capacity of the host to mount an appropriate antiviral defense aimed at limiting virus replication and spreading is critical to determine the outcome of the infection. As a consequence, the inability of the host to sustain an antiviral response leads to failure in eradicating the infection. Conversely, uncontrolled duration or intensity of the response is harmful to the host and is associated with the development of virus-associated pathogenesis, chronic inflammatory diseases and various autoimmune diseases ^5,6^. Therefore, these responses need to reach the ideal intensity and duration for efficient fighting of the infection while limiting tissue damage and promote tissue repair ^7^. To this aim, the various components of the host antiviral defense, including the transcriptional induction of interferons (IFNs) and proinflammatory cytokines and chemokines, but also the activation of cell death pathways, such as apoptosis, are subjected to stringent regulation by both positive and negative mechanisms ^8,9^.

Upon virus entry and sensing by the cytosolic pathogen recognition receptors (PRRs) retinoic acid-inducible gene I (RIG-I) and melanoma differentiation-associated protein 5 (MDA-5), the mitochondrial membrane-associated adaptor (MAVS) coordinates multiple signalling pathways ultimately leading to the activation of the transcription factors IFN regulatory factors (IRF) 3/7, Nuclear Factor κB (NF-κB) and Activator Protein 1 (AP-1) ^10-12^. Activation of IRF3 relies on a complex set of phosphorylations mainly mediated by the TANK-binding kinase 1 (TBK1)/ IκB kinase epsilon (IKKɛ) kinases ^13-17^. NF-κB, mainly p65/p50, activation during SeV and RSV infections occurs through IκB kinase (IKK)-dependent phosphorylation of the NF-κB inhibitor protein IκBα and of the p65 subunit ^18,19^. The signaling cascade leading to AP-1 activation is more elusive, but ultimately results in the phosphorylation of the nuclear Activating Transcription Factor 2 (ATF-2) and c-Jun subunits by JNK and p38 Mitogen-Activated Protein Kinases (MAPKs) ^10,20,21^. The activation of these transcription factors promotes the transcription of early antiviral genes, type I/III IFNs and proinflammatory cytokines and chemokines ^12,22,23^. In response to IFNs, hundreds of interferon-stimulated genes (ISGs) are induced to limit virus replication through enhancement of virus detection and innate immune signaling, cytoskeleton remodelling, inhibition of protein translation, induction of apoptosis, amongst other antiviral functions ^24-26^. These same PRRs have also been shown to activate IFN-independent cell death pathways, including apoptosis ^8^.

Post-translational modifications (PTMs), including phosphorylation, ubiquitylation and acetylation, of signal transduction proteins involved in the pathways engaged downstream of virus sensing are crucial to reach a fine-tuned regulation of the innate immune response ^27^. In light of the very well documented importance of phosphorylation PTMs in the regulation of the antiviral response, the protein phosphatases negatively regulating the signaling events have started to be identified. The exact role of the Ser/Thr protein phosphatase 1 (PP1) in the antiviral response remains elusive as PP1α and PP1γ were found to dephosphorylate MDA-5 and RIG-I leading to their activation ^28^, while they were also described to be responsible for the dephosphorylation of key C-terminal phosphoresidues of IRF3 leading to its inhibition ^29^. Most likely reflecting the complexity of IRF3 regulation through phosphorylation at multiple C-terminal phosphoacceptor sites, two other phosphatases, the Ser/Thr protein phosphatase 2 (PP2A) and MAPK phosphatase (MKP) 5, dephosphorylate IRF3 to terminate its activation ^30-32^. The Ser/Thr protein phosphatase, Mg^2+^/Mn^2+^-dependent (PPM) 1B acts as a TBK1 phosphatase to inhibit IRF3 activation, while PPM1A targets both TBK1/IKKɛ and MAVS for dephosphorylation to negatively regulate the antiviral response ^33,34^.

In the present study, we sought to address the role of the MKP-1/DUSP1 dual phosphatase (referred to thereafter as DUSP1) in the regulation of the host-defense against RSV and Sendai virus (SeV), a model paramyxovirus that is currently evaluated as a replication-competent backbone for the development of an RSV vaccine ^3^. We demonstrate that DUSP1 is a negative regulator of RSV- and SeV-induced JNK/p38 MAPK phosphorylation. However, this function is neither linked to the inhibition of the antiviral response nor to the induction of a cytokine and chemokine response elicited during virus infection. Interestingly, we found that interaction of JNK by the JNK interacting protein (JIP) 1 scaffold protein, previously shown to be critical for AP-1 and downstream cytokine production specifically, protects JNK from dephosphorylation by DUSP1. Although we confirmed that a JNK/p38 signalling module is involved in the induction of virus-induced apoptosis, our data suggests that DUSP1 has a pro-apoptotic function independently of JNK and p38 during SeV infection. Finally, we found that DUSP1 dampens cell migration during RSV infection. Altogether, these findings point to a previously unrecognized role of DUSP1 in functions that have an impact on virus-associated tissue damage and repair.

## RESULTS

### Induction of DUSP1 during SeV and RSV infection

To first assess a potential function of DUSP1, we analyzed DUSP1 expression during the course of SeV and RSV infection in A549 cells by immunoblot. DUSP1 was induced by about 4 fold at 6 h of RSV infection and was returned to basal levels at 24 h post-infection (**Figure 1A**). DUSP1 steady state levels remained unchanged during SeV infection (**Figure 1A**). Because DUSP1 turnover was previously reported to be affected by the ubiquitin-proteasome pathway ^35^, we further analyzed DUSP1 levels during SeV and RSV infection in the presence of the 26S proteasome inhibitor MG132. In the presence of MG132, induction of DUSP1 is detectable in both infections and the overall DUSP1 levels were markedly increased compared to control cells (**Figure 1B**). These results imply that DUSP1 protein levels are induced in response to SeV and RSV infection and that DUSP1 is subjected to proteasome-mediated degradation.

**Figure 1:**
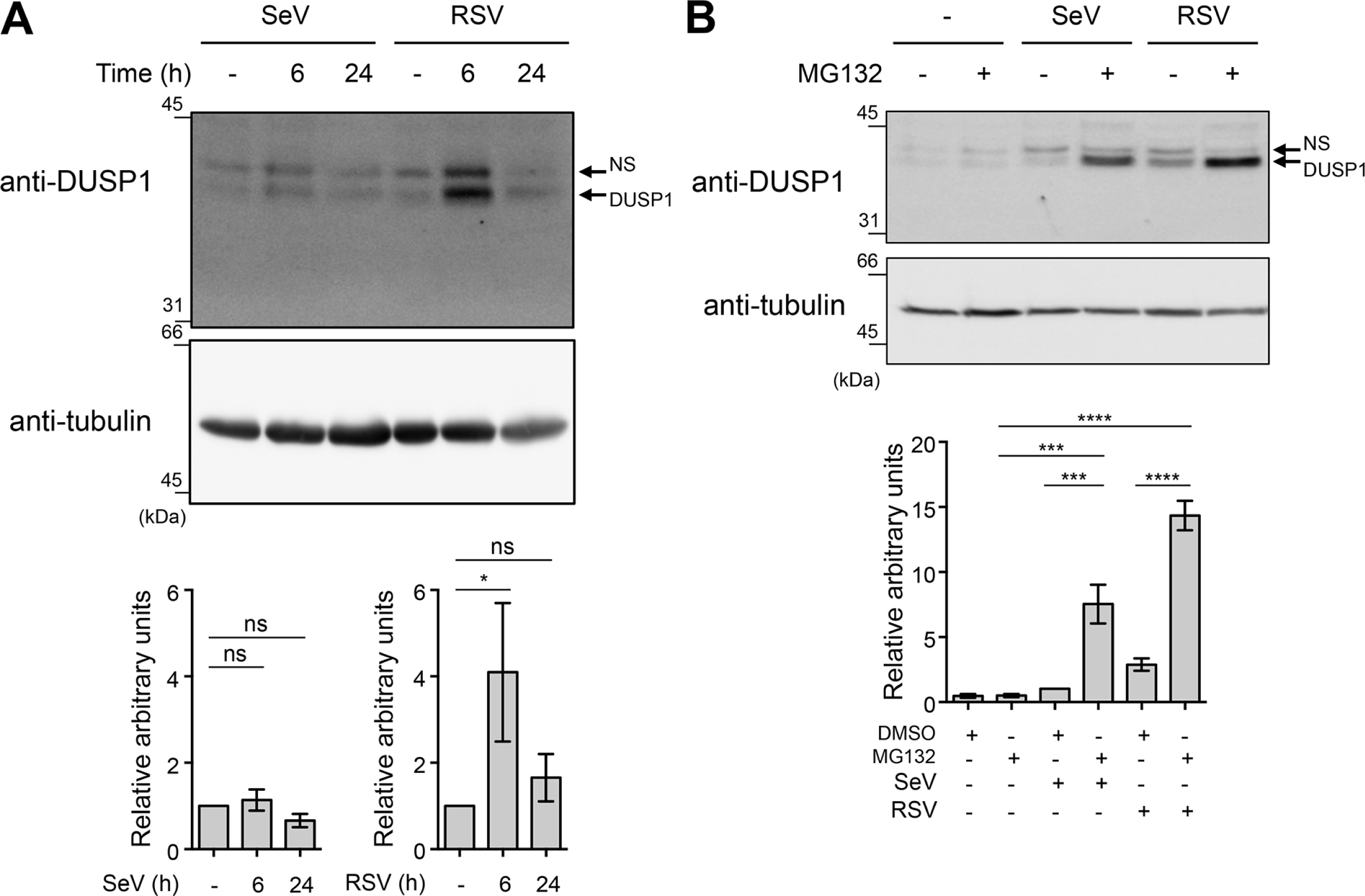
DUSP1 is induced during SeV and RSV infections and subjected to proteasomal degradation. In **A and B,** A549 cells were infected with SeV at 40 HAU/10^6^ cells or RSV at a MOI of 3 for the indicated times. **In B,** cells were pre-treated with MG132 (5 μM) or DMSO (vehicle) before infection for 6 h. WCE were analyzed by immunoblot using anti-DUSP1 specific antibodies. Detection of tubulin was used to control for equal loading. The data are representative of at least three independent experiments. Samples that are compared derive from the same experiment. Quantification of DUSP1 levels normalized over tubulin are represented as mean +/− SEM, n=7 **(A)** and n=3 **(B).** Statistical comparisons were performed using RM one-way ANOVA with Dunnett's **(A)** or Tukey's **(B)** post-tests. Full-length blots are presented in **Supplementary Figure 3**. NS: non-specific.

### DUSP1 does not affect SeV and RSV replication

Next, we sought to determine whether DUSP1 belongs to the numerous negative regulators of the antiviral response induced in response to virus infections ^9,36^. First, because of their importance in the control of the IFN-mediated antiviral defense, we evaluated whether DUSP1 has an impact on the signaling cascades leading to NF-κB and IRF3 activation. Considering previous characterization of NF-κB activation in the context of RSV infection ^37^, IκBα-S32 and p65-S536 phosphorylation were analyzed. Activation of IRF3 was monitored through detection of IRF3-S396 phosphorylation ^38^. Ectopic expression of DUSP1 in A549 cells did not alter IκBα, p65 or IRF3 phosphorylation profiles observed during the course of SeV (**Figure 2A and B**) or RSV infection (**Figure 2C**), showing that DUSP1 does not play a role in the regulation of these defense pathways. Additionally, ectopic expression of DUSP1 in A549 cells neither significantly altered SeV N mRNA levels measured by qRT-PCR (**Figure 2D top panel**) nor the *de novo* production of RSV virions quantified by plaque assay (**Figure 2D bottom panel**). Consistently, RNAi-mediated silencing of DUSP1 did not alter the capacity of RSV to replicate in A549 cells (**Figure 2E**). Together these data indicate that DUSP1 does not negatively regulate the capacity of the cell to mount an antiviral response to restrict RSV or SeV replication.

**Figure 2:**
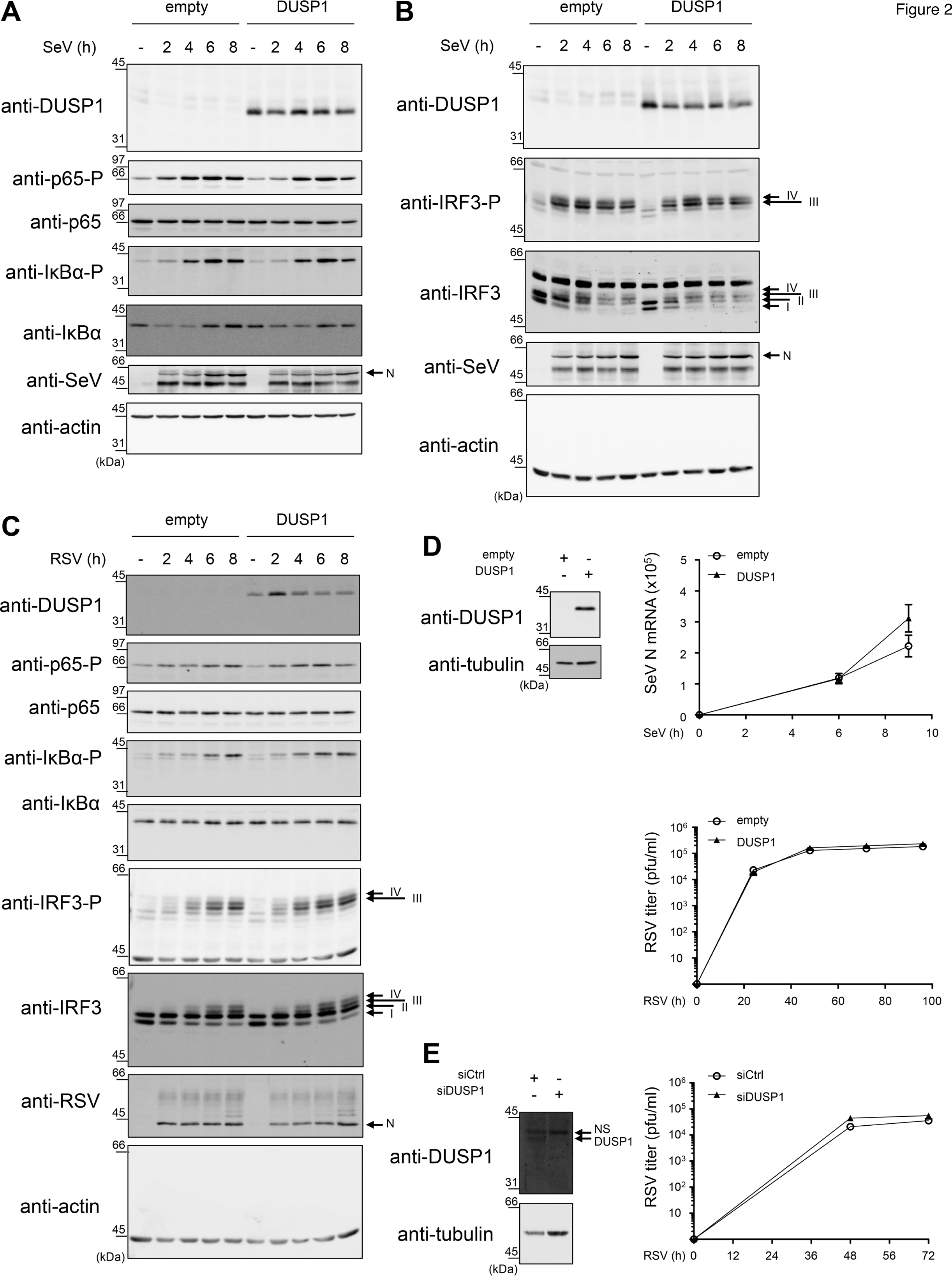
DUSP1 expression does not alter the antiviral pathways triggered by SeV and RSV infections. A549 cells were transfected with an empty- or DUSP1-expressing plasmid **(A-D)** or with Ctrl or DUSP1 specific siRNA **(E)** before infection with SeV at 40 **(A and B)** or 5 HAU/10^6^ cells (**D, top panel**) or with RSV at MOI of 3 **(C)** or 1 **(D, bottom panel and E)** for the indicated times. In **A-C,** immunoblot analyses were performed to detect DUSP1, phosphorylated p65 (p65-P), total p65, phosphorylated IκBα (IκBα-P), total IκBα, phosphorylated IRF3 (IRF3-P) and total IRF3. Infection was monitored using anti-SeV or anti-RSV antibodies (N proteins are shown). Detection of actin was used as a loading control. The data are representative of three different experiments. Samples that are compared derive from the same experiment. Full-length blots are presented in **Supplementary Figure 4**. **In D,** SeV N mRNA was quantified by qRT-PCR. In **D and E,** the release of infectious RSV virions was quantified by plaque forming unit (pfu) assay. Data are represented as mean +/− SEM, n=6 **(D, top panel)** or n=8 **(D, bottom panel and E)** and analyzed using two-way ANOVA with Bonferroni's post-test.

### DUSP1 inhibits RSV- and SeV-induced JNK and p38 phosphorylation

Because DUSP1 preferentially dephosphorylates JNK and p38 in a variety of inflammatory contexts ^39,40^, we next sought to evaluate the role of DUSP1 in the negative regulation of RSV- and SeV-induced JNK and p38 activation. First, we monitored the effect of ectopically expressed DUSP1 on the SeV- and RSV-induced JNK T183/Y185 and p38 T180/Y182 phosphorylation in A549 cells. As expected, immunoblot analysis revealed that SeV (**Figure 3A and B**), and to a lesser extent RSV (**Figure 3C**), induce JNK and p38 phosphorylation. Importantly, DUSP1 expression abrogates JNK and p38 phosphorylation (**Figure 3A-C**). Subsequently, we confirmed the role of DUSP1 through analysis of the impact of DUSP1 silencing on the profile of virus-induced JNK and p38 phosphorylation. Immunoblot analysis confirmed the efficiency of DUSP1 silencing in A549 cells throughout the course of SeV and RSV infection (**Figure 4**). Consistent with a role of DUSP1 in the dephosphorylation of JNK and p38, we observed a marked increase of JNK and p38 phosphorylation in the absence of DUSP1 during SeV (**Figure 4A**) and RSV infection (**Figure 4B**). Altogether, these observations reveal a critical role of DUSP1 in the negative regulation of JNK and p38 activation during RSV and SeV infection.

**Figure 3:**
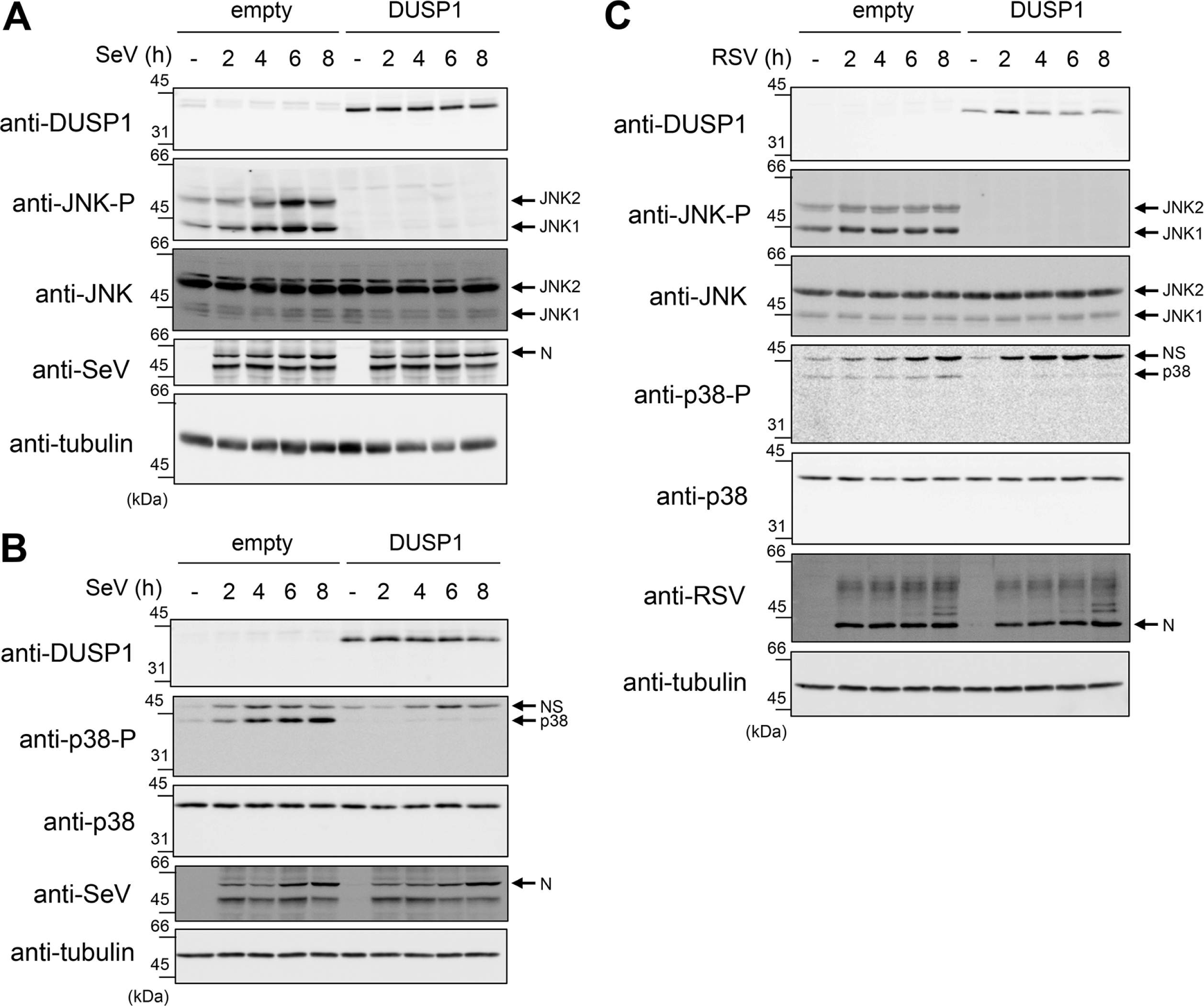
Ectopic expression of DUSP1 inhibits JNK and p38 phosphorylation elicited by SeV and RSV infections. A549 cells were transfected with an empty- or DUSP1-encoding plasmid before infection with SeV at 40 HAU/10^6^ cells **(A and B)** or RSV at a MOI of 3 **(C)** for the indicated times. Levels of DUSP1, phosphorylated JNK (JNK-P), total JNK, phosphorylated p38 (p38-P), total p38, SeV N and RSV N protein levels were monitored by immunoblot. Equal loading was verified using anti-tubulin antibodies. The data are representative of at least three independent experiments. Samples that are compared derive from the same experiment and blots were processed in parallel. Full-length blots are presented in **Supplementary Figure 5**. NS: non-specific.

**Figure 4:**
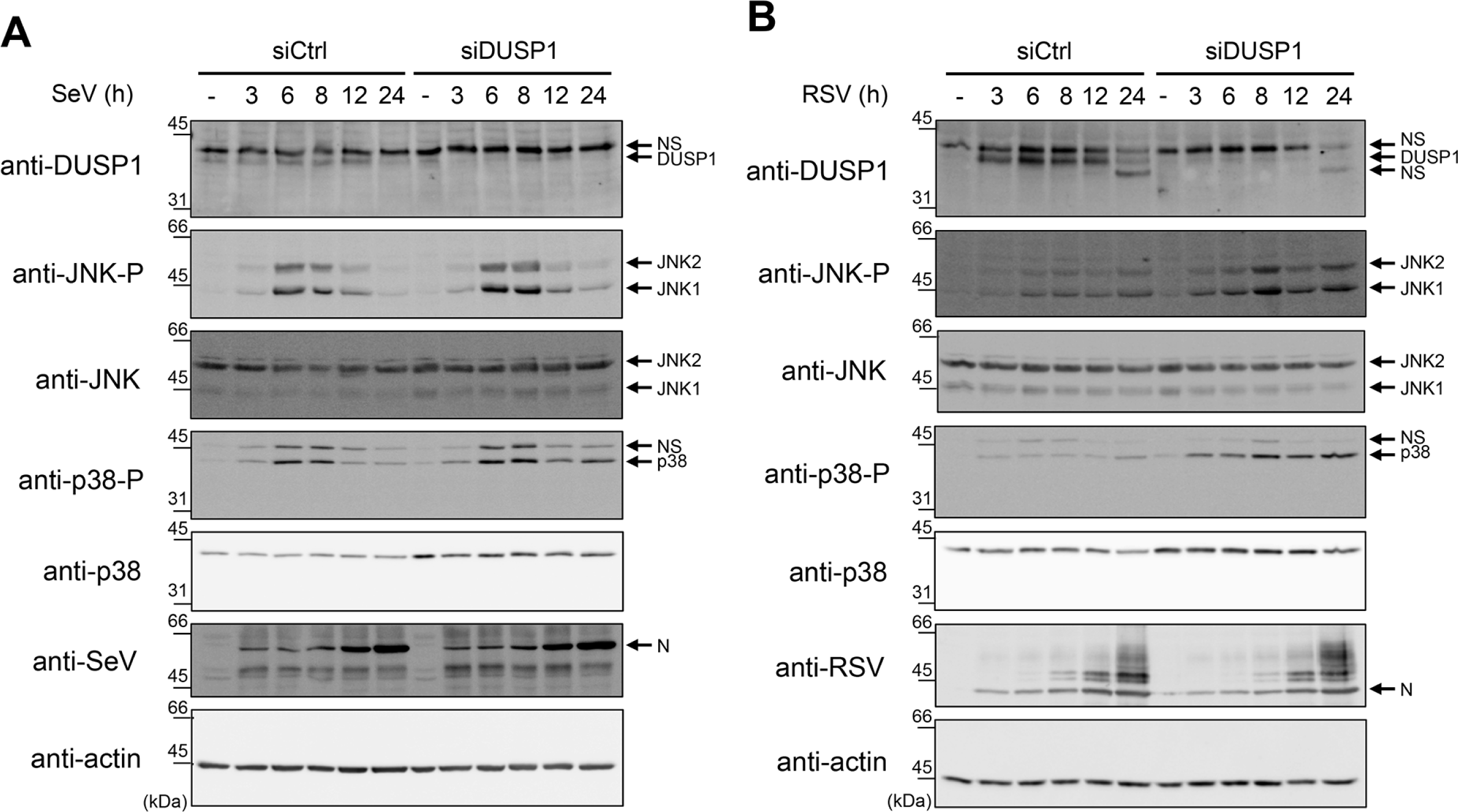
DUSP1 silencing increases SeV- and RSV-induced JNK and p38 phosphorylation. A549 cells were transfected with Ctrl or DUSP1-specific siRNA before being left uninfected or infected with SeV at 40 HAU/10^6^ cells **(A)** or RSV at MOI of 3 **(B)** for various times. Immunoblot analyses were performed to verify the efficiency of DUSP1 silencing and expression levels of phosphorylated JNK (JNK-P), total JNK, phosphorylated p38 (p38-P) and total p38. Infection was monitored using anti-SeV and anti-RSV antibodies (N proteins are shown). Detection of actin was used as a loading control. The data are representative of at least three independent experiments. Samples that are compared derive from the same experiment and blots were processed in parallel. Full-length blots are presented in **Supplementary Figure 6**. NS: non-specific.

### Interaction with JIP1 protects JNK from DUSP1 dephosphorylation

Scaffolding proteins of the JIP family interact with distinct pools of JNK to specifically place them next to their upstream activators and downstream substrate mediating specific functional pathways ^41,42^. Therefore, we aimed to investigate the role of JIP proteins in RSV- and SeV-induced JNK activation and regulation by DUSP1. The JIP family is composed of 4 members, JIP1-4, with distinct expression profiles. Here, we assessed the role of JIP1 and JIP3 in A549 cells based on previous reports indicative of their expression in lung cells ^43-46^. In contrary, JIP2 was shown to be restricted to human brain ^47^ and human JIP4 was found only in testis ^48,49^. First, we tested the interaction between JNK and JIP1 or JIP3 during SeV and RSV infection. Ectopically expressed JIP1 coimmunoprecipitated with endogenous JNK, both in uninfected and SeV- and RSV-infected cells (**Figure 5A**). By contrast, JIP3 did not interact with JNK (**Figure 5A**). Since we demonstrated the existence of a JIP1/JNK complex, we examined the role of JIP1 in the regulation of DUSP1 dephosphorylation during SeV and RSV infection. A549 cells were cotransfected with JIP1 together with empty or DUSP1-expressing constructs before infection with SeV (**Figure 5B**) or RSV (**Figure 5C**). Confirming our finding (**Figure 3**), ectopic expression of DUSP1 resulted in inhibition of SeV- and RSV-induced JNK phosphorylation (**Figure 5B and C**). Importantly, when JIP1 was coexpressed with DUSP1, the levels of phosphorylated JNK was increased 5.9 and 4 times compared to cells transfected with DUSP1 only during SeV and RSV infection, respectively (**Figure 5D**). Collectively, these data demonstrate that JIP1 interacts with JNK and that recruitment of JNK by JIP1 prevents dephosphorylation by DUSP1 during SeV and RSV infection.

**Figure 5:**
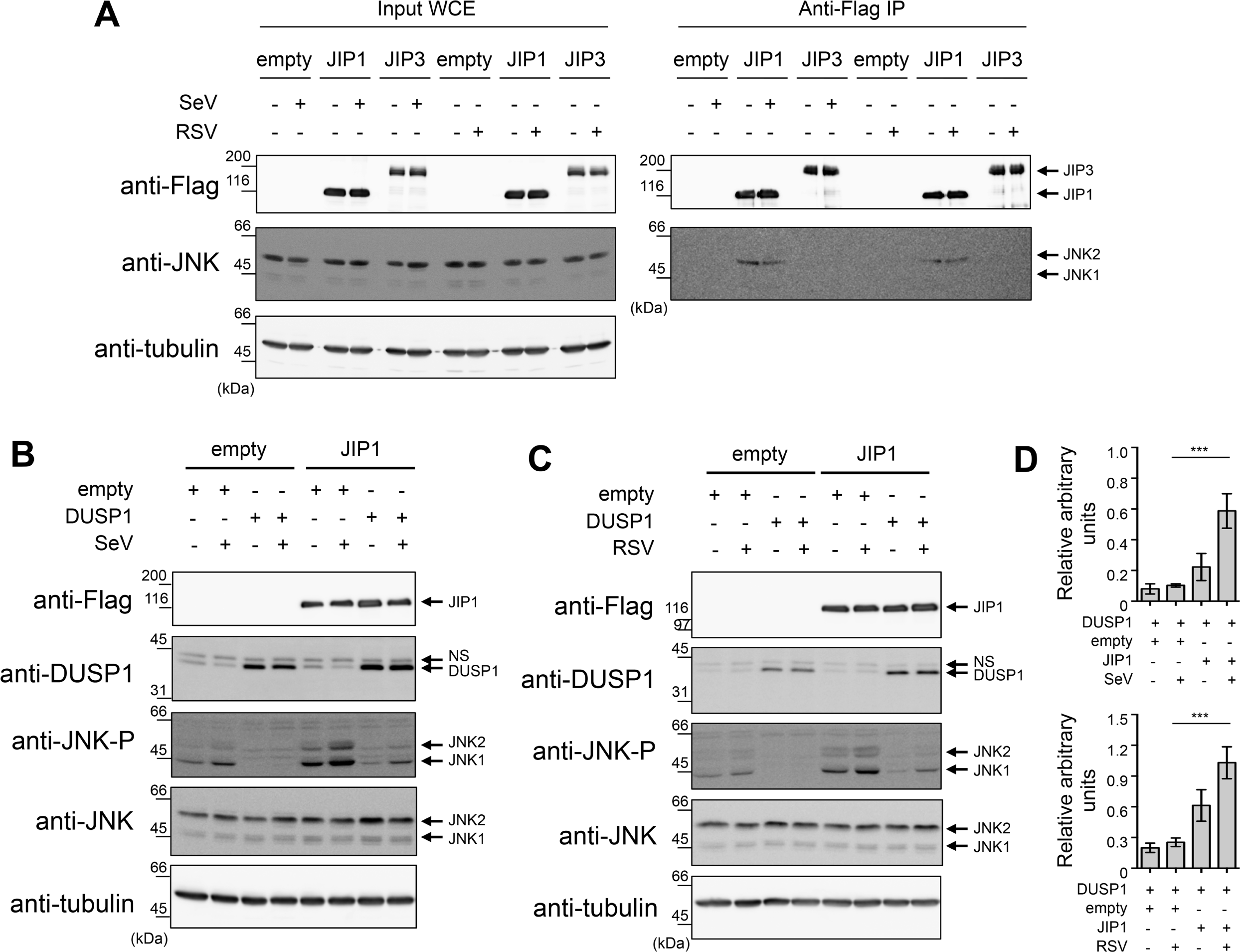
JIP1 scaffold interacts with JNK and protects from dephosphorylation by DUSP1. A549 cells were transfected with the indicated combination of Flag-JIP1-, Flag-JIP3- and DUSP1-expressing plasmids or the corresponding empty plasmids, and left uninfected or infected with SeV at 40 HAU/10^6^ cells or RSV at MOI of 3 for 6 h. **In A,** WCE were subjected to immunoprecipitation (IP) using anti-Flag antibodies followed by immunoblot. In **B and C,** WCE were resolved by SDS-PAGE before immunoblot. In **A, B and C,** levels of protein expression were assessed using anti-Flag, anti-phosphorylated JNK (JNK-P), anti-JNK and anti-DUSP1 antibodies. Tubulin was used as a loading control. Data are representative of at least three different experiments. Samples that are compared derive from the same experiment. **In D,** quantification of phosphorylated JNK levels normalized over total JNK in DUSP1 expressing cells in the absence or presence of JIP1 are shown. Mean +/− SEM, n≥3. Statistical comparisons were performed using RM one-way ANOVA with Tukey's post-test. Full-length blots are presented in **Supplementary Figure 7**. NS: non-specific.

### AP-1 and downstream cytokine production elicited during infection are protected from DUSP1-mediated inhibition of JNK and p38

JIP1 is essential for the activation of the JNK/c-Jun axis and downstream AP-1-dependent cytokine expression ^50-53^. A main function of JNK and p38 in the host defense against virus infection is to regulate the activation of AP-1 (ATF-2/c-Jun) that participates in the enhanceosome structure controlling IFNβ expression and contributes to the regulation of proinflammatory cytokine expression ^10,20,21^. The observation that JIP1 prevents JNK from being dephosphorylated by DUSP1 suggests a model in which AP-1-dependent functions would be protected from the negative regulation of DUSP1 during the infection. To confirm that JNK/p38 were involved in virus-induced AP-1 activation and downstream cytokine expression, A549 cells were infected with SeV in the absence or presence of JNK (SP600125) and p38 (SB203580) inhibitors. Immunoblot analysis showed that inhibition of JNK and p38 significantly decreased the levels of phosphorylated c-Jun and ATF-2 (**Figure 6A**). Accordingly, inhibition of JNK and p38 significantly impaired the induction of IFNβ, CCL2, CXCL8 and IL-6 by SeV, quantified using Luminex-based assays (**Figure 6D and Supplemental Figure 1A**). These results confirmed the role of the JNK/p38-AP-1 signaling cascade in the regulation of SeV-mediated cytokine induction and therefore prompted us to evaluate the impact of ectopic expression of DUSP1 on this pathway. Although ectopic expression of DUSP1 prevents JNK and p38 phosphorylation, SeV- and RSV-induced phosphorylation of c-Jun and ATF-2 remained similar to control cells (**Figure 6B and C**). Analysis of the expression profile of a panel of 48 cytokines using Luminex-based multiplex assays during SeV infection revealed that none of them were significantly altered following DUSP1 ectopic expression (**Figure 6E and Supplemental Figure 1B**). Additionally, silencing of DUSP1 did not affect the levels of IFNβ, CCL2, CCL5, CXCL8 and IL-6 induced by SeV (**Figure 6F and Supplemental Figure 1C)**. Measurement of SeV-induced *IFNB, CXCL8* and *CCL2* mRNA levels revealed no impact of DUSP1 ectopic expression, excluding a role of DUSP1 in the transcriptional regulation of cytokines (**Supplemental Figure 2**). Similar results were observed during RSV infection, with silencing of DUSP1 having no impact on the induction of IFNβ, CCL2, CCL5, CXCL8 and IL-6 (**Figure 6G**). Altogether these observations point to a selective role of DUSP1 in the negative regulation of JNK and p38 during SeV and RSV infection, leaving the AP-1 pathway and downstream cytokine production intact.

**Figure 6:**
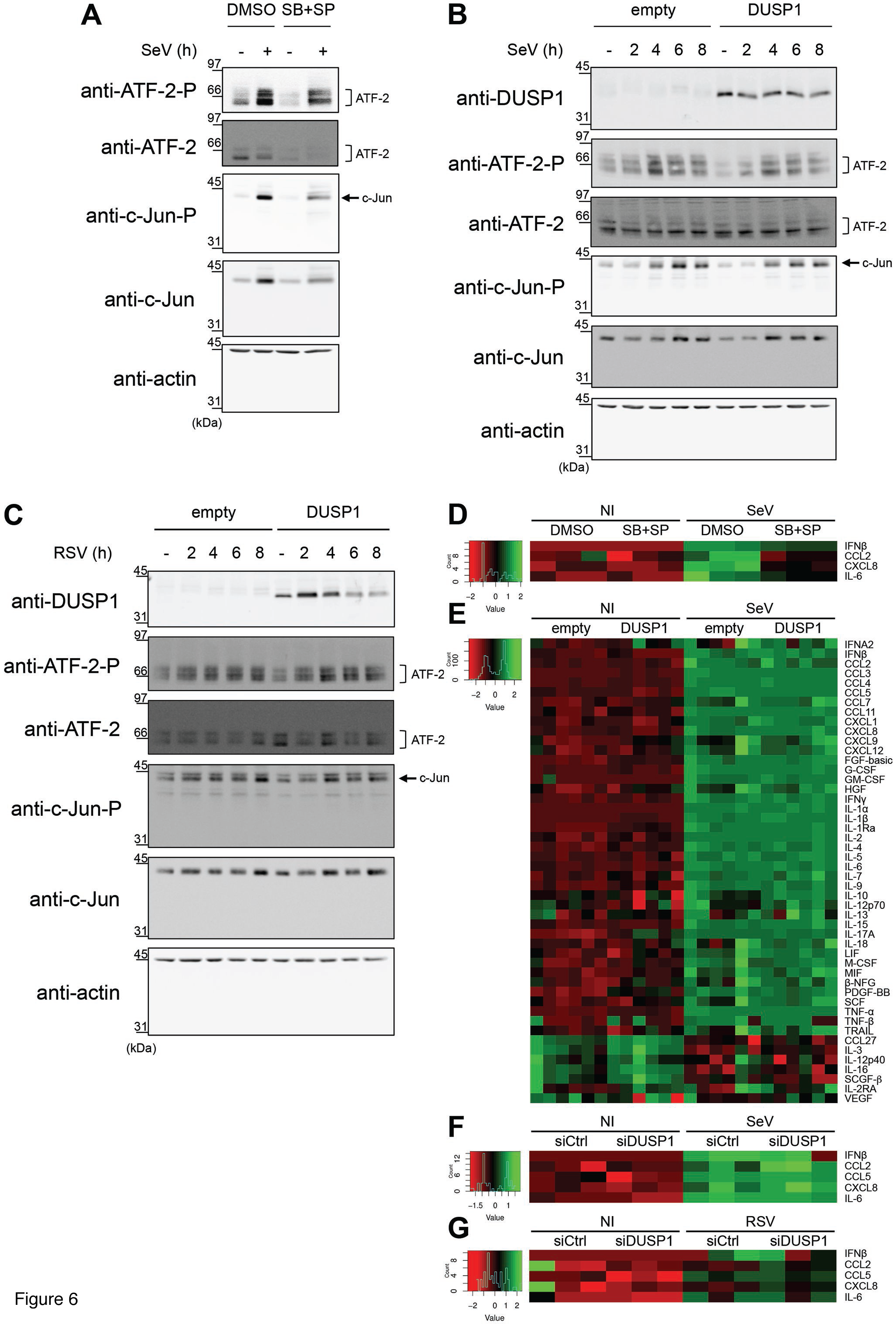
DUSP1-mediated inhibition of JNK and p38 leaves virus-mediated activation of AP-1 and cytokine production intact. A549 cells were either pretreated with DMSO (vehicle) or SB203580 (10 μM) + SP600125 (10 μM) for 30 min prior to infection **(A and D)**, transfected with empty or DUSP1-expressing plasmids **(B, C and E)** or transfected with Control (siCtrl) or DUSP1-specific siRNA **(F and G)** before infection with SeV at 40 HAU/10^6^ cells **(A, B, D, E and F)** or RSV at MOI of 3 **(C and G)** for the indicated times or for 6 h **(A and D)**, 16 h **(E)** or 12 h **(F and G)**. In **A-C,** levels of phosphorylated ATF-2 (ATF-2-P), total ATF-2, phosphorylated c-Jun (c-Jun-P), total c-Jun and DUSP1 were analyzed by immunoblot. Actin was used to verify equal loading. The data are representative of three independent experiments. Samples that are compared derive from the same experiment and blots were processed in parallel. Full-length blots are presented in **Supplementary Figure 8**. In **D-G,** release of cytokines was quantified using Luminex-based multiplex assays. Heatmaps represent cytokine levels (pg/ml) logarithmically transformed, centered and scaled, measured in each biological replicates (n=3 in **D, F** and **G**, n=6 in **E**). Scatter plots of cytokines levels are shown in **Supplemental Figure 1**.

### DUSP1 promotes SeV-induced apoptosis independently of JNK/p38 inhibition

JNK/p38 MAPK pathways are also known to play a critical role in the regulation of apoptosis, being either pro- or anti-apoptotic depending on the context ^54-59^. A MAVS/JNK pathway has been shown to be indispensable for SeV to induce apoptosis ^60,61^. Therefore, we sought to assess the impact of DUSP1 on the JNK/p38-dependent apoptosis triggered by SeV. A549 cells were transfected with siCtrl or siDUSP1 followed by treatment with DMSO (vehicle) or SP600125 and SB203580 to inhibit JNK and p38 before SeV infection. Quantification of Annexin V+ cells, corresponding to early and late apoptosis, was performed by Flow cytometry analysis of Annexin V/PI stained cells **(Figure 7A)**. Confirming a moderate role of JNK/p38 in SeV-induced apoptosis, inhibition of JNK/p38 reduced the number of Annexin V+ cells by 12 %. Unexpectedly, DUSP1 silencing also similarly decreased SeV-induced apoptosis by 10 %. Importantly, when DUSP1 silencing and JNK/p38 inhibition were combined, SeV-induced apoptosis was decreased by 23 %, showing an additive effect of the two pathways. SeV-induced apoptosis involves the intrinsic pathway leading to caspase 9 and 3 activation ^62^. Therefore, to confirm the impact of JNK/p38 inhibition and/or DUSP1 silencing on SeV-induced apoptosis, caspase 9 and 3 cleavage was monitored by immunoblot. Confirming the results of Annexin V, both inhibition of JNK and p38 and silencing of DUSP1 significantly impaired SeV-induced caspase 3 cleavage (**Figure 7B**). Additionally, caspase 9 cleavage that was induced following SeV infection confirming the engagement of the intrinsic pathway, was reduced in the absence of DUSP1 compared to control cells (**Figure 7B)**. These findings confirmed that a JNK/p38 signaling module is involved in the induction of apoptosis during SeV infection, but they also argue against a negative regulation of this pathway by DUSP1. Rather, the observations support the hypothesis that DUSP1 is required for induction of the intrinsic apoptotic pathway independently of JNK and p38.

**Figure 7:**
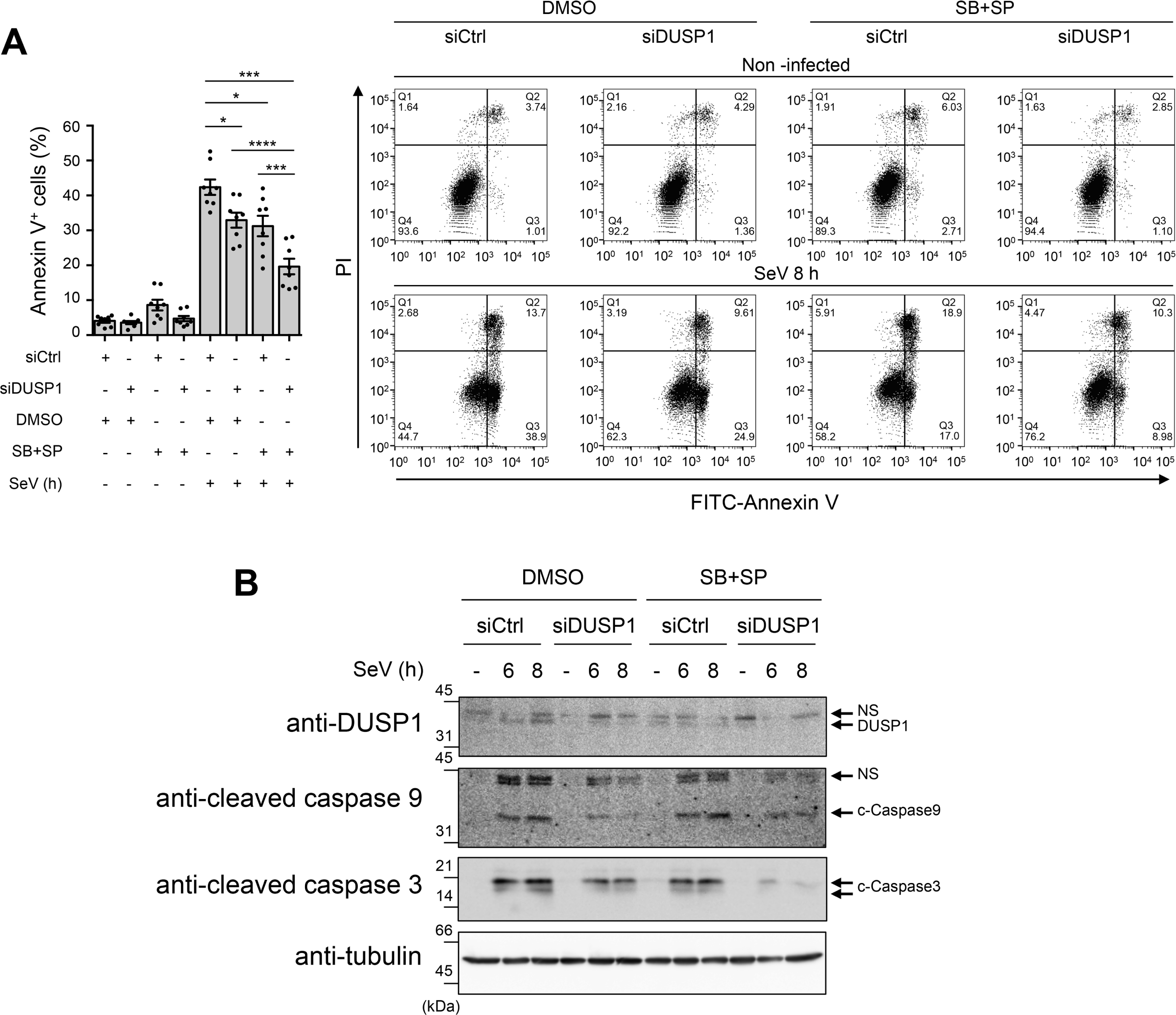
DUSP1 and JNK/p38 act on independent SeV-induced pro-apoptotic pathways. A549 cells were transfected with control (siCtrl) or DUSP1-specific siRNA for 48 h and pretreated with DMSO (vehicle) or SB203580 + SP600125 (10 μM each) for 30 min before infection with SeV at 40 HAU/10^6^ cells for 8 h **(A)** or the indicated times **(B)**. **In A,** cells were harvested and stained with Annexin V-FITC and PI and analyzed by flow cytometry. The left bar graph represents the percentage of Annexin V positive (Annexin V^+^/PI^−^ and Annexin V^+^/PI^+^; Q2 and Q3) apoptotic cells. Mean +/− SEM, n=8 independent replicates. Statistical comparisons amongst infected cells were performed by RM one-way ANOVA with Tukey's post-test. Representative FACS plots are shown. **In B,** DUSP1, cleaved Caspase 9 (c-Caspase 9) and cleaved Caspase 3 (c-Caspase 3) protein levels were monitored by immunoblot. Equal loading was verified using anti-tubulin antibodies. The data are representative of three independent experiments. Samples that are compared derive from the same experiment and blots were processed in parallel. Full-length blots are presented in **Supplementary Figure 9**. NS: non-specific.

### DUSP1 negatively regulates cell migration during RSV infection

In contrast to SeV, RSV does not induce significant apoptosis at early time of infection (^63^ and data not shown) when DUSP1 dephosphorylates JNK/p38. This prompted us to evaluate the role of DUSP1 is alternative functions during RSV infection. Compelling evidence has implicated JNK/p38 in the positive regulation of cell migration in several cell types ^64^. Phenotypic observation of A549 cells during RSV infection following DUSP1 silencing showed obvious formation of pseudopods that were not observed in control cells (**Supplementary Movie 1**). This pointed to a propensity of infected cells to stretch out and migrate in the absence of DUSP1. To investigate the role of DUSP1 in cell migration during RSV infection, 2D cell migration dynamics was assessed by single-cell tracking using time-lapse video-microscopy in subconfluent A549 cell cultures either left uninfected or infected with RSV following siCtrl or siDUSP1 transfection (**Supplementary Movie 1**). As hypothesized, cell trajectories and migration rates were altered following DUSP1 silencing. In the absence of DUSP1, RSV-infected cells were migrating further from their origin compared to control cells that rather adopted a trajectory in circles around their point of origin (**Figure 8**). These observations demonstrate a critical negative role of DUSP1 in the regulation of cell migration during RSV infection.

**Figure 8:**
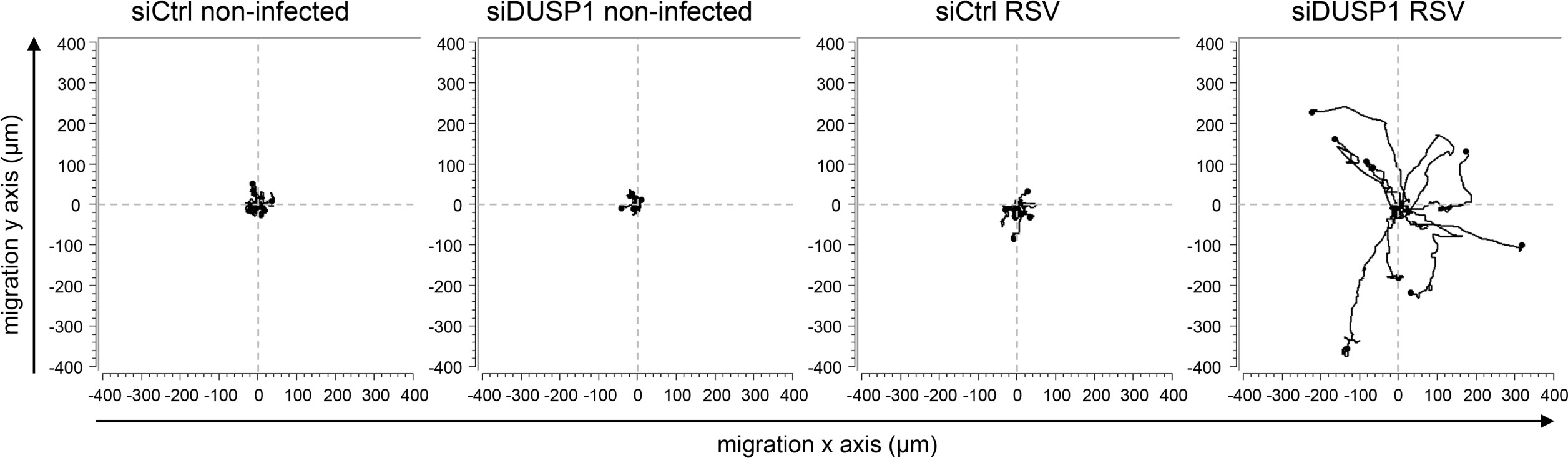
DUSP1 impairs cell migration during RSV infection. A549 cells were transfected with control (siCtrl) or DUSP1-specific siRNA before RSV infection at a MOI of 3 for 8 h. Single-cell tracking was performed using live-video microscopy to monitor two-dimensional cell migration. Trajectories of 9 cells per conditions are presented. The origins of cell trajectories have been aligned to start at the position (x = 0; y = 0). The data are representative of two independent experiments for non-infected conditions and four independent experiments for RSV-infected conditions. Original live video imaging of cell tracking are available in **Supplementary Movie 1**.

## DISCUSSION

DUSP1 is expressed in various cell types and tissues and mainly acts as a critical negative regulator shaping the duration, magnitude and spatiotemporal profile of p38 and JNK MAPKs, and to a lesser extent ERK, activation following physiological and pathological stimuli ^65^. As such, DUSP1 has been extensively shown to negatively regulate the innate immune anti-bacterial defense and the cellular response to allergens ^66,67^. In these contexts, DUSP1 mainly inhibits proinflammatory cytokine expression ^40,68-73^. The role of DUSP1 in the innate response to virus infection is far less known and, to our knowledge, was not previously assessed in the context of infection by RSV or SeV.

In the present study, we show that DUSP1 negatively regulates p38 and JNK phosphorylation induced by SeV and RSV (**Figure 3 and 4**). Analysis of virus replication, of the NF-κB and IRF3 signaling pathways and of the induction of IFNs levels excluded a role of DUSP1 in the negative regulation of the antiviral defense (**Figure 2 and 6**). The observation that pharmacological inhibition of JNK/p38 MAPKs during SeV infection significantly decreased downstream ATF-2/c-Jun activation and cytokine induction (**Figure 6A and D**) confirmed the previously documented existence of a JNK-p38/AP-1 signaling axis critical for virus-induced cytokine production ^10,20,21^. Unexpectedly, DUSP1-mediated inhibition of JNK/p38 phosphorylation had no effect on downstream phosphorylation of ATF-2/c-Jun or on the levels of a wide panel of cytokines elicited during infection (**Figure 6**). We hypothesized that this observation might reflect the existence of different subsets of JNK segregated through the interaction with specific scaffold proteins. In the MAPK signaling modules involving JNK, members of the JIP family of scaffold proteins selectively enhance specific JNK-dependent functions by interacting with and linking the upstream kinases to JNK activation ^41,42^. In our experimental model, we showed that JNK interacts with JIP1, but not with JIP3, at basal levels and during infection (**Figure 5A**). Importantly, we found that the JIP1/JNK interaction dampens the capacity of DUSP1 to dephosphorylate JNK (**Figure 5B-D**). Compelling evidence demonstrates that JIP1 is essential for the activation of the JNK/AP-1 axis that controls cytokine expression ^50-53^. Although we cannot exclude other mechanisms, it is reasonable to speculate that during SeV and RSV infection, the sequestration of JNK through molecular interaction with JIP1 protects JNK and downstream AP-1 and cytokine response from DUSP1 negative regulation (**Figure 6**). Further studies will be required to fully address this model. Previous reports have shown that interaction with JIP1 leads to the retention of JNK in the cytoplasm ^43,74^. DUSP1 is a nuclear phosphatase ^75,76^ and therefore predominantly dephosphorylates JNK and p38 in the nucleus ^77^. It is thus very likely that retention of JNK in the cytoplasm by JIP1 contributes to protect JNK from DUSP1 dephosphorylation. Our results differ from previous reports showing DUSP1-dependent inhibition of proinflammatory cytokine expression following challenge with the viral dsRNA mimetic poly I:C, coronavirus and vaccinia virus infection ^71,78,79^. Additionally, DUSP1 was shown to enhance vaccinia virus replication ^79^. Thus, DUSP1 appears to differentially regulate the cytokine response and virus replication depending on the virus. One might speculate that the JIP1/JNK interaction might be differently affected upon infection by distinct viruses. While we did not observe changes of the JIP1/JNK complex during SeV and RSV infection, other viruses may interfere with this interaction thereby making JNK available for dephosphorylation by DUSP1 to negatively regulate downstream cytokine release. Although to date no virus was reported to dissociate the JIP1/JNK complex, it is interesting to note that Vaccinia virus-encoded B1R kinase interacts with JIP1 leading to increased binding of JNK to JIP1 and downstream activation of c-Jun ^80^.

Amongst alternative functions driven by JNK and p38 MAPK pathways are the regulation of cell proliferation and apoptosis ^81^. Cell cycle analysis during SeV and RSV infection following ectopic expression or silencing of DUSP1 failed to demonstrate a role of DUSP1 in cell proliferation (data not shown). Instead, we found that DUSP1 is required for induction of the intrinsic apoptotic pathway triggered during SeV infection (**Figure 7B**). Confirming previous reports ^60,82,83^, we also found that a JNK/p38 pathway contributes to SeV-induced apoptosis (**Figure 7A**). The observation that inhibitions of DUSP1 and JNK/p38 have additive effect on the negative regulation of SeV-induced apoptosis (**Figure 7**) strongly hints at a model in which DUSP1 regulates virus-induced apoptosis in a JNK/p38-independent manner. Further studies will be required to challenge this model. Additionally, this also argues toward a model in which the JNK/p38 pro-apoptotic function is also protected from DUSP1 dephosphorylation, possibly through the observed JNK/JIP1 interaction. This model is consistent with the fact that MAVS-MKK7-JNK defines a pro-apoptotic pathway during SeV infection and that JIP1 specifically functions in the JNK pathway by tethering MLK3 (MAPKKK), MKK7 (MAPKK) and JNK ^60,74^. Moreover, the JIP1/JNK axis has also been shown to be pro-apoptotic in neurons exposed to stress ^51,84^. RSV does not induce significant apoptosis at early time of infection (^63^ and data not shown) when DUSP1 dephosphorylates JNK/p38, but DUSP1 silencing resulted in formation of pseudopods that were not observed in controls cells **Supplementary Movie 1**). In the quest to characterize DUSP1-dependent JNK and p38 functions, we thus assessed the impact of DUSP1 on cell migration during RSV infection. Indeed, accumulating evidence implicates JNK and p38 in pro-migratory functions in various contexts ^64^. Here, we demonstrate that in the absence of DUSP1, cell migration of RSV infected cells was highly enhanced (**Figure 8**) pointing to a critical role of DUSP1 in the negative regulation of JNK/p38-mediated cell migration during RSV infection. In contrary to RSV, SeV induces high levels of apoptosis at early time of infection, which prevented an unbiased analysis of the migration parameters. Regulation of apoptosis and cell migration are part of the arsenal of the host response that have the potential to not only influence the outcome of virus spreading, but also the extent of virus-induced tissue damage and repair thereby contributing to pathogenesis ^7,85^. The implication of the observed DUSP1-mediated regulation of apoptosis and cell migration in SeV and RSV spreading can be excluded based on the lack of effect of DUSP1 ectopic expression or silencing on virus replication (**Figure 2**). Induction of apoptosis and inhibition of cell migration by DUSP1 rather suggests a role in the induction of cell damage and inhibition of tissue repair and thereby in virus-induced pathogenesis.

Altogether our results suggest a model in which DUSP1 is a negative regulator of important host mechanisms that limit tissue damage and promote tissue repair. An exaggerated cytokine response can also contribute to host self-inflicted damages and thereby contribute to virus-induced pathogenesis ^86^. We found that the cytokine response remains intact upon manipulation of DUSP1 expression due to protection by the interaction of JNK with JIP1 (**Figure 5A**). The observation that JIP1/JNK interaction prevents a potential negative regulation of the cytokine response by DUSP1 opens avenues for specific therapeutic targeting. Indeed, one can hypothesize that inhibition of the JIP1/JNK interaction might restore DUSP1-dependent inhibition of the AP-1/cytokine axis during the infection, while leaving other JIP1-independent JNK functions unaffected. Further studies will be required to evaluate this possibility. The JIP1/JNK interaction has long been considered an interesting specific target and this led to the development of small molecules and peptides that inhibit the interaction between JNK and JIP1 and efficiently block JNK activity toward selective substrates, including ATF-2 and c-Jun ^87-90^. Alternatively, direct inhibition of DUSP1 ^91,92^ might be a strategy to improve tissue homeostasis, by reducing virus-induced apoptosis and restoring cell migration, during virus infection.

## MATERIALS AND METHODS

### Cell culture

A549 cells (American Type Culture Collection, ATCC) were grown in Ham F12 medium (GIBCO) and Vero cells (ATCC) in Dulbecco's Modified Eagle Medium (DMEM, GIBCO). Both media were supplemented with 10 % heat-inactivated fetal bovine serum (HI-FBS, GIBCO) and 1 % L-Glutamine (GIBCO). Cultures were performed without antibiotics and were tested negative for mycoplasma contamination (MycoAlert Mycoplasma Detection Kit, Lonza) every 2 months.

### Infections

Subconfluent monolayers (90 % confluency) of A549 cells were infected with Sendai virus (SeV, Cantell strain, Charles River Laboratories) at 5-40 hemagglutinin units (HAU)/10^6^ cells as indicated in serum free medium (SFM). At 2 h post-infection, the medium was supplemented with 10 % HI-FBS and the infection pursued for the indicated time. Infection with RSV A2 (Advanced Biotechnologies Inc), amplified and purified as described in ^37^, was performed at a MOI of 1-3 as indicated in medium containing 2 % HI-FBS. Where indicated, A549 were pretreated with 5 μM MG132 (Calbiochem) or DMSO (vehicle, Sigma-Aldrich) for 1 h before infection. Pretreatment with 10 μM SB203580 and 10 μM SP600125 (Invivogen) or the corresponding vehicle DMSO was performed for 30 min prior to infection.

### SDS-PAGE and immunoblot

The procedures used for preparation of Whole Cell Extracts (WCE), resolution by SDS-PAGE electrophoresis and immunoblot was fully detailed in ^93^. The following primary antibodies were used in this study: anti-MKP-1/DUSP1 (M18, #sc-1102), anti-α-tubulin (B-7, #sc-5286) and anti-NFκB p65 (C-20, #sc-372) were obtained from Santa Cruz Biotechnology. Anti-phosphoT183/Y185 SAPK/JNK (G9, #9255), anti-SAPK/JNK (#9252), anti-phosphoT180/Y182 p38 MAPK (D3F9, #4511), anti-p38 MAPK (D13E1, #8690), anti-phosphoS536 NF-kappaB p65 (93H1, #3033), anti-phosphoS32 IκBα (14D4, #2859), anti-IκBα (#9242), anti-phosphoT71 ATF-2 (11G2, #5112), anti-ATF-2 (20F1, #9226), anti-phosphoS73 c-Jun (D47G9, #3270), anti-c-Jun (60A8, #9165), anti-cleaved Caspase 9 (D315, #9505) and anti-cleaved Caspase 3 (D175, #9664) were from Cell Signaling. Anti-actin clone AC-15 (#A5441) and anti-Flag M2 (#F1804) were purchased from Sigma-Aldrich, anti-IRF3 (#39033) was from Active Motif and anti-RSV (#AB1128) was from Chemicon International. Anti-phosphoS396 IRF3 was described in ^38^ and anti-SeV was obtained from Dr. J. Hiscott, McGill University, Montreal, Canada. HRP-conjugated goat anti-rabbit and rabbit anti-goat (Jackson Laboratories), and goat anti-mouse (Kirkegaard & Perry Laboratories) were used as secondary antibodies. Immunoblots were quantified using the ImageQuantTL software (Molecular Dynamics).

### Plasmids

The human DUSP1-pCMV6XL5 plasmid was obtained from Origene. The pcdna3 Flag JIP1b (Addgene plasmid # 52123) and pcdna3 Flag JIP3b (Addgene plasmid # 53458) plasmids were a gift from Roger Davis ^50,94^. All constructs were verified using Sanger sequencing at the McGill University and Génome Québec Innovation Centre, Montréal, Canada.

### Plasmid transfection

Plasmid transfection in A549 cells was achieved using the TransIT-LT1 transfection reagent (Mirus). Briefly, a total of 3 μg of DNA was transfected per 35 mm plates of A549 cells at 70 % confluency using a transfection reagent/DNA ratio of 1:2. Transfection was pursued for 24 h to 48 h before further treatment, as indicated.

### qRT-PCR

Total RNA were extracted using the RNAqueous-96 Isolation Kit (Ambion) and quantified. Reverse transcription was performed using 1 μg total RNA using the Quantitect reverse Transcription kit (Qiagen). Specific mRNA levels were quantified by qPCR using Fast start SYBR Green Kit (Roche) for *SeV N* (S: agtatgggaggaccacagaatgg, AS: ccttcaccaacacaatccagacc). A reaction without RT and a reaction with H_2_O were performed with each run to ensure absence of genomic DNA contamination. Fluorescence was collected using the Rotor-Gene 3000 Real Time Thermal Cycler (Corbett Research). Results were analyzed by the ΔΔCT method after normalization to S9 mRNA levels (S: cgtctcgaccaagagctga, AS: ggtccttctcatcaagcgtc).

### Virus titration by plaque forming unit (pfu) assay

RSV infectious virions were quantified by methylcellulose plaque forming units assay. Briefly, supernatant from infected plates was harvested and subjected to serial dilutions before being used to infect monolayers of Vero cells. After 2 h of infection, Vero cells were washed and covered with 1 % methylcellulose prepared in DMEM containing 2 % FBS. RSV plaques were immunodetected at 5 days post-infection. After removal of the methylcellulose, cells were fixed in 80 % methanol for 30 min and air-dried. Plates were incubated for 15 min at room temperature (RT) in PBS 0.1X pH 7.2 containing 2 % milk and 0.05 % tween before being incubated with anti-RSV antibodies (#AB1128, Chemicon International) for 3 h at RT, washed 3 times in PBS 0.1 X pH 7.2 containing 0.05 % tween and finally incubated with rabbit anti-goat antibodies (Jackson Laboratories) for 1 h at RT. After 3 more washes, plates were incubated with Enhanced Chemiluminescence substrate (ECL, Perkin Elmer Life Sciences) for 5 min at RT and chemiluminescence signal was detected using a CCD camera-based apparatus (LAS4000 mini, GE healthcare). Quantification of plaques was performed using the ImageQuantTL software (Molecular Dynamics) and expressed as pfu/ml.

### RNAi transfection

RNAi oligonucleotide (ON-target siRNA, Dharmacon) transfection was performed using the Oligofectamine reagent (Invitrogen). A non-targeting sequence, described in ^13^, was used as control. For efficient DUSP1 expression silencing, a mix of two siRNA sequences, caguuauggugaugacuuauu and ccgacgacacauauacauauu, were used. Cells were plated at 30 % confluency and transfected for 24-48 h before further treatment depending on specific experimental design.

### Co-immunoprecipitation experiments

WCE were prepared as described in the immunoblot section. For co-immunoprecipitation studies, 1-1.5 mg of WCE were incubated with 2 μg anti-Flag M2 antibodies (#F1804, Sigma-Aldrich) linked to 30 μl protein G Sepharose beads (Sigma-Aldrich) for 3 h at 4 °C. Beads were washed 5 times with lysis buffer and the elution of immunocomplexes was performed by incubation with a solution of 100 μg/ml Flag peptide (#F3290, Sigma-Aldrich) prepared in lysis buffer for 2 h at 4 °C. Immunoprecipitates were then denatured with 1:4 (v/v) 5X loading buffer ^93^ and heated for 5 min at 100 °C. WCE and immunoprecipitates were resolved by SDS-PAGE as described in the immunoblot section.

### Luminex-based quantification of cytokines

Supernatants from infected A549 cells were collected and centrifuged for 5 min at 1000 g followed by a 10 min centrifugation at 10 000 g to remove cell debris. RSV-infected supernatants were UV-inactivated for 20 min before further analysis. Cytokines were quantified by Luminex xMAP Technology using the Bio-Plex Pro*^TM^* Human Cytokine Standard Plex Group I and Group II or custom assembled plex for IFNβ, IL-6, CXCL8, CCL2 or CCL5 from Bio-Rad on a Bio-Plex 200 System (Bio-Rad). Analyses were performed using the Bio-Plex Manager Software version 6.0.0.617. Heatmaps were produced from the raw expression analysis data using *R* ^95^ and the *gplots* package ^96^ on logarithmically transformed, centered and scaled data.

### Detection of apoptosis by flow cytometry (Annexin V/PI staining)

For detection of apoptosis, the supernatant of cell culture was collected and cells were harvested using 0.05 % trypsin-EDTA. Supernatant and cells were pooled and centrifuged for 5 min at 250 g at 4 °C. The cell pellet was resuspended in Annexin V buffer (10 mM Hepes pH 7.4, 150 mM NaCl, 5 mM KCl, 1 mM MgCl_2_ and 1.8 mM CaCl_2_) and stained using 1 μl of FITC-Annexin V (#556419, BD Biosciences) and 1 μl of Propidium Iodide (PI, 1 mg/ml, #P-4170, Sigma-Aldrich) per 2-5 × 10^5^ cells for 10 min at 4 °C. Immediately after staining, acquisition of fluorescence was performed on a BD LSR II flow cytometer (BD Biosciences) using the BD FACSDiva Software (BD Biosciences). A minimum of 10 000 cells was analyzed for each sample. Data were analyzed using the FlowJo software (FlowJo, LLC).

### Analysis of cell migration

Two-dimensional cell migration of single cells (9 cells/picture and 4 pictures/condition) was evaluated by single-cell tracking using live-video microscopy. Images were captured at 5 min intervals over an 18 h period by digital camera connected to a Zeiss microscope (Axio Observer.Z1) as in ^97^. The cell trajectories were analysed using the Image J software (National Institutes of Health, Bethesda, MD, USA).

### Statistical analyses

Quantitative results are represented as mean +/− SEM. Statistical comparisons were performed with Prism 7 software (GraphPad) using the indicated tests. The following *P*-values were considered significant: P<0.05 (*), P<0.01 (**), P<0.001 (***) or P<0.0001 (****).

## AUTHOR CONTRIBUTIONS

ACR, NZ, DA, EB and NG conceived and designed the experiments. ACR, EC, NZ, EM, DA, MM, AG and AF performed experiments. ACR, NZ, DA and NG analyzed the data. ACR and NG wrote the manuscript. All co-authors edited and approved the manuscript.

## ACKNOWLEDGEMENTS

We thank members of the laboratory and of the immunovirology group at CRCHUM for fruitful discussions. We also thank François-Christophe Marois-Blanchet and François Harvey at the CRCHUM bioinformatics facility and Dr. Dominique Gauchat and Annie Gosselin at the CRCHUM flow cytometry platform.

The present work was funded by grants from the Canadian Institutes of Health Research (CIHR) [MOP-130527] and by the Research Chair in signaling in virus infection and oncogenesis from the Université de Montréal to NG. ACR was supported by graduate studentships from CIHR, the Fonds de la Recherche du Québec en Santé (FRQS) and the Réseau en Santé Respiratoire du FRQS. AG was recipient of a studentship from the Natural Sciences and Engineering Research Council of Canada (NSERC). NZ was recipient of a post-doctoral fellowship from FRQS/INSERM (Québec/France) and DA by a post-doctoral fellowship from FRQS. NG was recipient of a Tier II Canada Research Chair.

